# A probabilistic view of protein stability, conformational specificity, and design

**DOI:** 10.1101/2022.12.28.521825

**Authors:** Jacob A. Stern, Tyler J. Free, Kimberlee L. Stern, Spencer Gardiner, Nicholas A. Dalley, Bradley C. Bundy, Joshua L. Price, David Wingate, Dennis Della Corte

**Author notes:** Correspondence to: Dennis Della Corte < >.

## Abstract

Several recently introduced approaches use neural networks as probabilistic models for protein sequence design. These models use various objective functions and optimization schemes. The choice of objective function and optimization scheme comes with trade-offs that are not always well explained. We introduce probabilistic definitions of protein stability and conformational specificity and show how these chemical properties relate to the *p*(structure| seq) objective used in recent protein design algorithms. This links probabilistic objective functions to experimentally testable outcomes. We present a new sequence decoding algorithm, termed “BayesDesign”, that uses Bayes’ Rule to maximize the *p*(structure| seq) objective. We evaluate BayesDesign in the context of two protein model systems, the NanoLuc enzyme and the WW structural motif.

## 1. Introduction

A common protein sequence design method is to find the minimum-energy sequence for a desired protein structure (Norn et al., 2021). This approach has been used since the first Rosetta energy function was introduced (Simons et al., 1997) (Liu & Kuhlman, 2006), continuing into current usage with improved energy functions (Alford et al., 2017). Over the years, a number of algorithms have been developed to find the minimum energy sequence for a given structure (Jones, 1994) (Dahiyat & Mayo, 1997) (Kuhlman et al., 2003) (Ingraham et al., 2019) (Anand et al., 2022).

However, finding the minimum-energy sequence for a given structure does not guarantee that the designed sequence folds into the desired structure. As observed by (Norn et al., 2021), minimizing the absolute energy of the sequence for the structure can result in sequences with energy minima at other conformations. To address this, (Norn et al., 2021) uses gradient descent over sequence space to maximize *p*(structure|seq), such that predicted structure matches target structure, using trRosetta (Yang et al., 2020) as a model for *p*(structure|seq).

Several subsequent works attempt to maximize the improved, *p*(structure| seq) objective. ColabDesign (Roney & Ovchinnikov, 2022) (Wang et al., 2022) uses a input optimization method similar to (Norn et al., 2021) but with AlphaFold (Jumper et al., 2021) as the structure prediction model. Similarly, (Anishchenko et al., 2021) uses tr-Rosetta as a model for *p*(structure|seq) but uses Markov Chain Monte Carlo (MCMC) optimization over sequences to maximize the KL divergence between *p*(structure| seq) for a designed sequence and *p*(structure seq) for a “background” structure.

However, the authors of both (Norn et al., 2021) and (Anishchenko et al., 2021), found that optimizing over the inputs of a forward structure prediction model leads to “adversarial” sequences that, while predicted to fold to the target structure, are insoluble in practice (Dauparas et al., 2022) (Wang et al., 2022). This is a known problem exploited in “adversarial optimization”, where optimizing over the inputs to a model finds inputs for which the model has high confidence, but which do not lie on the manifold of data for which the model makes reliable predictions (Goodfellow et al., 2014).

There have been several attempts to resolve the problem of insoluble adversarial sequences. (Norn et al., 2021) and (Anishchenko et al., 2021) introduce terms the original objective to increase the probability that sequences fall on the manifold of training data for the structure prediction model. (Dauparas et al., 2022) attempts to resolve the adversarial problem by using ProteinMPNN (which is trained with a *p*(seq |structure) objective) to redesign protein sequences for backbones designed by (Anishchenko et al., 2021). But all of these solutions introduce alter the original design objective such that designed sequences are different from the sequences that maximize *p*(structure|seq).

Two common protein design goals are stability and conformational specificity (Marshall & Mayo, 2001). Stability describes the energy difference between the folded native state and the unfolded denatured state of a protein. Conformational specificity refers to how strongly a protein prefers one folded conformation over other folded conformations.

In this work, we present formal probabilistic definitions of protein stability and conformational specificity which illustrate the relationship between these protein design criteria and the *p*(structure = *X*| seq) objective. This provides a link between a probabilistic objective function and experimentally testable properties. The link to protein stability and conformational specificity motivates careful adherence to the *p*(structure = *X*| seq) objective function without introducing other terms that change its theoretical properties.

Next, we describe a new approach to designing sequences maximizing the *p*(structure|seq) objective without relying on gradient descent or MCMC optimization techniques that lead to adversarial sequences. This approach uses existing models for *p*(seq|structure) and *p*(seq) and applies Bayes’ Rule to design proteins that maximize *p*(structure|seq).

Finally, we evaluate the stability and conformational specificity of BayesDesign-designed proteins on two model systems: the luminescent NanoLuciferase (NanoLuc) enzyme and the WW beta sheet motif.

Our work introduces probabilistic definitions of two important properties of a protein. It introduces “BayesDesign”, a method to maximize the *p*(structure|seq) objective while avoiding the pathologies involved in optimizing over inputs to a neural network.

Our contributions are as follows:

- We mathematically formalize protein design objectives for protein stability and conformational specificity and show how they relate to the Boltzmann probability objective function *p*(structure|seq).
- We derive a tractable probabilistic model, “BayesDesign” to design protein sequences maximizing the Boltzmann probability objective *p*(structure|seq) without finding adversarial sequences.
- We provide experimental evaluation of the stability and conformational specificity of protein sequences designed by the BayesDesign algorithm.

## 2. Theory

### 2.1. Formalizing protein design objectives

#### 2.1.1. Boltzmann probability

The **Boltzmann probability** of a protein conformation *X* refers to the probability that a protein will be in that conformational state:

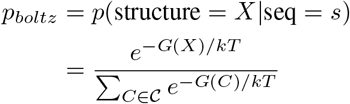

where 𝒞 is the set of all possible states and *G*(*C*) refers to the Gibbs free energy of a state *C*. This is the objective function described in (Norn et al., 2021). The denominator can be partitioned into three terms:

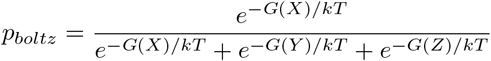

where *Y* is the set of alternate folded conformations, and *Z* is the set of denatured conformations. The rationale for partitioning the denominator into these three terms is given in section 2.1.4.

For notational simplicity throughout this paper, we write *G*(*X*) to refer to the Gibbs free energy of a structure *X* conditioned on its corresponding sequence *s*.

#### 2.1.2. Protein stability

**Protein stability** is defined as the Gibbs free energy difference between the denatured state and native state:

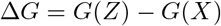

where *X* is the minimum-energy folded state and *Z* is the denatured state.

Maximizing protein stability is equivalent to maximizing the probability ratio between the minimum-energy folded state *X* and the denatured state *Z* (see Proposition 1 in the SI):

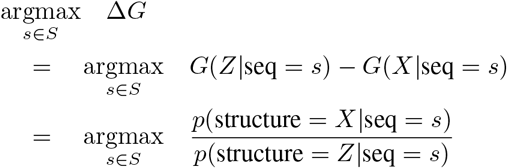

where *G*(*X*|seq = *s*) is the Gibbs free energy of the native state for a given sequence and *G*(*Z*| seq = *s*) is the Gibbs free energy of the denatured state for a given sequence *s*.

Based on this equivalence, we propose a **probabilistic definition of protein stability** that is not equal, but which is maximized when the original definition of protein stability is maximized:

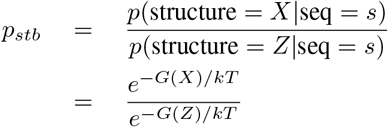

Writing the definition of protein stability in its probabilistic form, we can see the relationship between protein stability and the Boltzmann probability objective - they differ only in the terms present in the denominator.

#### 2.1.3. Conformational specificity

We also introduce a **probabilistic definition of conformational specificity** which serves as a compliment to the probabilistic definition of protein stability. A sequence that maximizes this metric maximizes the probability ratio between the native conformation and alternate folded conformations.

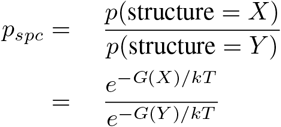

Conformational specificity measures the tendency of a protein structure to maintain one conformation over other folded conformations. This is important, as proteins with high conformational specificity to a are less likely to adopt another conformation that may aggregate (Zhu et al., 2015), driving the dynamic equilibrium of conformations toward the undesirable conformation.

Whereas protein stability compares the probability of the native folded state to the denatured state, conformational specificity compares the probability of the native state to alternate folded conformations.

#### 2.1.4. Relationship between Boltzmann probability, protein stability, and conformational specificity

If a sequence design approach assumes a model *p*(structure = *X*| seq) of the Boltzmann probability of a given conformation given a sequence, then maximizing that objective has predictable effects on protein stability and specificity.

These effects are made apparent by partitioning the denominator of the Boltzmann probability objective into three terms, as shown in section 2.1.1. There are three ways to maximize *p*(structure = *X*| seq): 1) by reducing the Gibbs free energy *G*(*X*) of the native state *X*, 2) by increasing the Gibbs free energy *G*(*Z*) of the denatured state *Z*, or 3) by increasing the Gibbs free energy *G*(*Y*) of alternate folded conformations.

Increasing *p*(structure = *X*|seq) by deepening the energy well of the structure *X* will increase both stability and conformational specificity. Increasing *p*(structure = *X*| seq) by increasing the energy of the denatured state *Z* will increase stability without affecting conformational specificity, and increasing *p*(structure = *X*| seq) by removing low-energy alternate conformations *y* ∈*Y* will increase conformational specificity without affecting stability.

Thus, assuming that the probability model is correct, maximizing *p*(structure = *X*|seq) will increase either the stability or conformational specificity of that state, and possibly both. However, it is possible that maximizing *p*(structure = *X*| seq) may increase stability at the cost of conformational specificity or vice versa, depending on which offers the greatest improvement to the Boltzmann probability.

Figure 2 shows a hypothetical case where the sequence that maximizes Boltzmann probability (Seq A) is different from the sequence that maximizes stability (Seq B) or conformational specificity (Seq C). Figure S1 illustrates a case (i.e. a set of energy landscapes) where optimizing Boltzmann probability would result in selecting the maximum stability sequence, and Figure S2 illustrates a case where optimizing Boltzmann probability would result in selecting the maximum specificity sequence.

**Figure 1.**
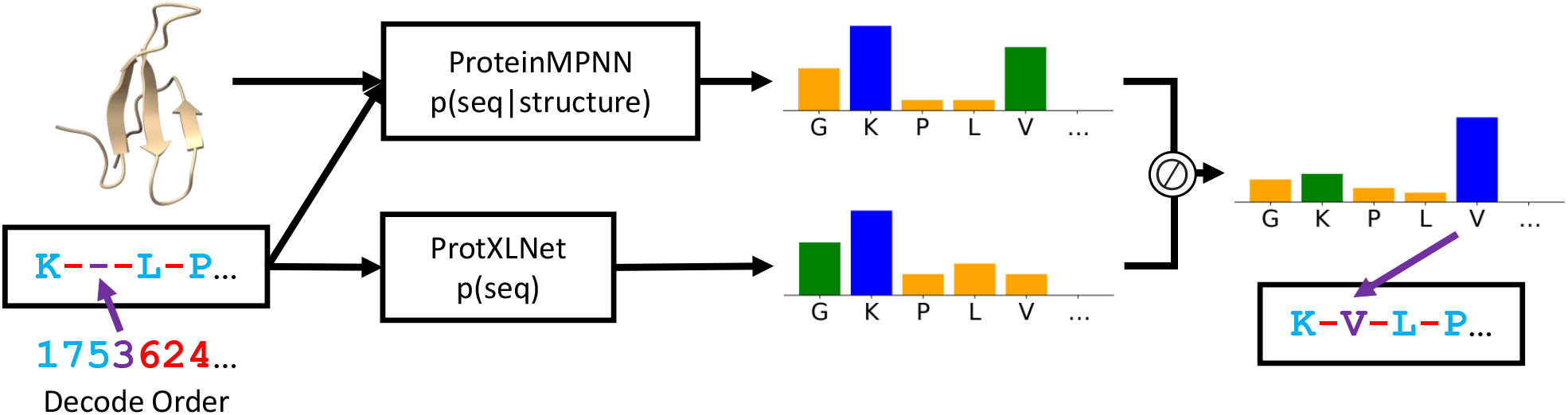
The BayesDesign model for de novo protein design.

**Figure 2.**
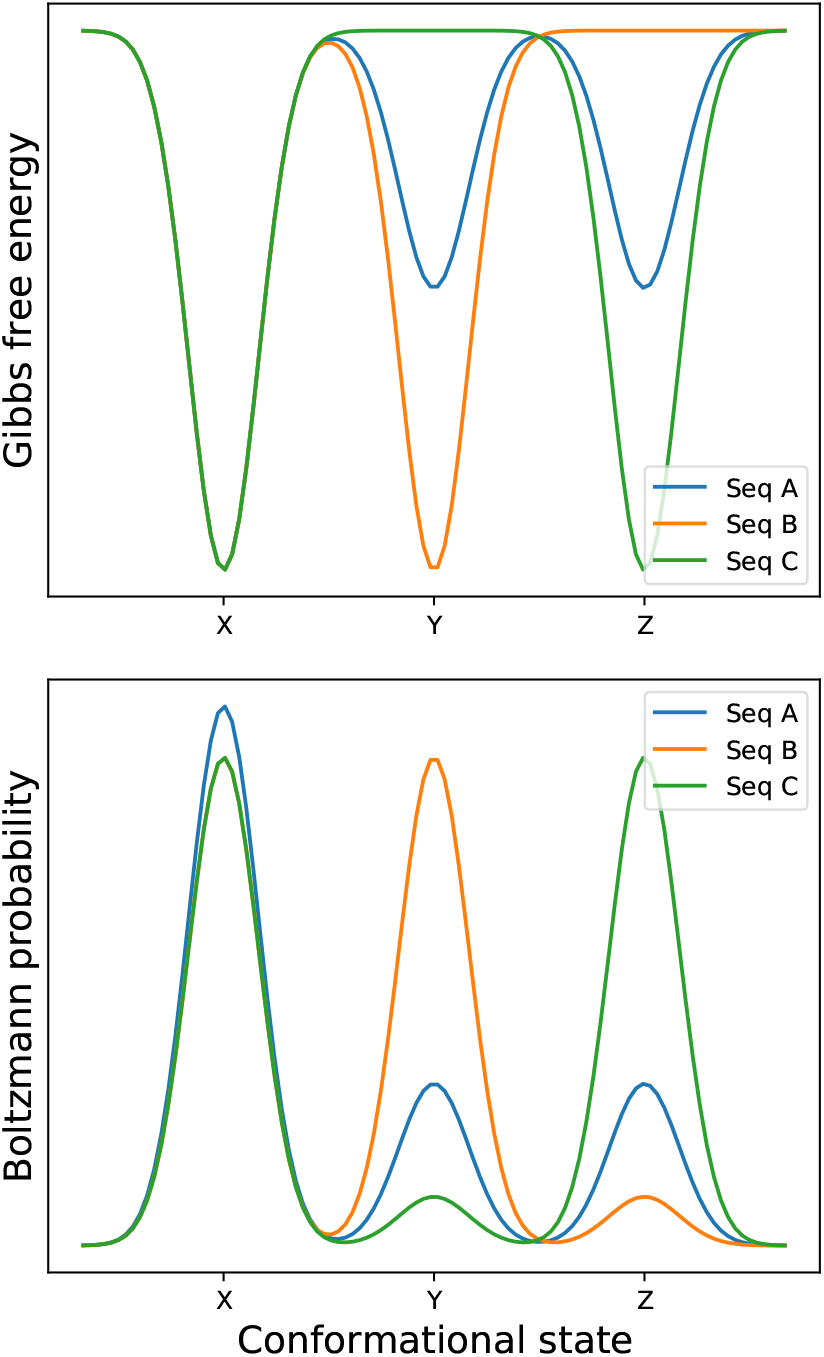
Protein sequences can have equal Gibbs free energy for a given structure, while having varying Boltzmann probability, stability, and conformational specificity, depending on the Gibbs free energy of other states. *X, Y*, and *Z* denote the native state, the alternate folded state, and the denatured state, respectively.

### 2.2. Current sequence design objectives

For comparison, we examine two sequence design objectives commonly used in the current literature.

#### 2.2.1. 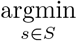 *G*(structure = *X*, seq = *s*)

This is the objective used when designing a sequence that minimizes the Rosetta energy function (Alford et al., 2017). Sequences designed via this objective often have rugged folding landscapes, and while the sequence is the minimum-energy sequence for the structure, the structure may not be the minimum-energy structure for the sequence (Norn et al., 2021).

#### 2.2.2 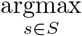 *p*(seq = *s*|structure = *X*) OR 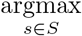 *p*(seq = *s*, structure = *X*)

First, we note that maximizing *p*(seq, structure = *X*) over sequences is equivalent to maximizing *p*(seq structure = *X*):

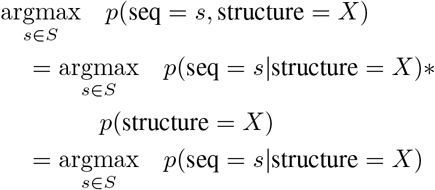

(Norn et al., 2021) discusses the importance of optimizing *p*(structure = *x*| seq), but the objective they use in practice is an objective that maximizes *p*(seq, structure = *X*), factorized into *p*(structure = *X*|seq) and *p*(seq). The added *p*(seq) term is intended to minimize the divergence of designed sequences from known sequences. However, it voids the theoretical implications of maximizing *p*(structure = *X*| seq) (namely, maximizing Boltzmann probability and by extension protein stability and/or conformational specificity).

(Anishchenko et al., 2021) maximizes a *p*(seq, structure) objective similar to (Norn et al., 2021), but does not condition on a structure *X*.

(Dauparas et al., 2022) maximizes *p*(seq| structure = *X*) directly, training an autoregressive model to predict protein sequence, conditioned on structure.

Similar to (Dauparas et al., 2022), several other state-of-the-art protein design algorithms involve a sequence decoder trained to maximize *p*(seq| structure) (Watson et al., 2022) (Ingraham et al., 2022).

### 2.3. Bayes’ Rule to maximize *p*(structure|seq)

We propose a new algorithm based on Bayes’ Rule to maximize the Boltzmann probability objective *p*(structure|seq).

#### 2.3.1. Probabilistic Model

Our goal is to design sequences maximizing *p*(structure|seq) without using gradient-based or MCMC optimization over the inputs of a deep neural network. To do this, we use Bayes’ Rule to rewrite the objective:

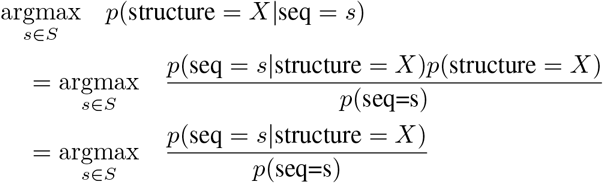

This means that we can maximize *p*(structure = *X*|seq) by using two models, *p*(seq |structure = *X*) and *p*(seq), and use a sequence decoding scheme instead of optimizing over sequences. This results in faster sequence design without the problem of adversarial optimization.

There are a few properties that would be especially useful for these two models. First, we want them to be autoregressive models of the joint probability. That allows us to use greedy algorithms like beam search to decode. It is also useful for them to be order-agnostic probabilistic models. This allows us to choose a decoding order that designs the most important sequence locations first (e.g. around a catalytic site).

We use ProteinMPNN (Dauparas et al., 2022) as a model for *p*(seq |structure) and ProtXLNet (Elnaggar et al., 2020) for *p*(seq). Both models possess the desired order-agnostic and autoregressive properties.

#### 2.3.2. Decoding

We can factorize the joint probability of a sequence as the product of conditional probabilities using the chain rule of probability. This reduces to a ratio of probabilities for each position in the sequence, where at each position the probability ratio gives a score for each amino acid:

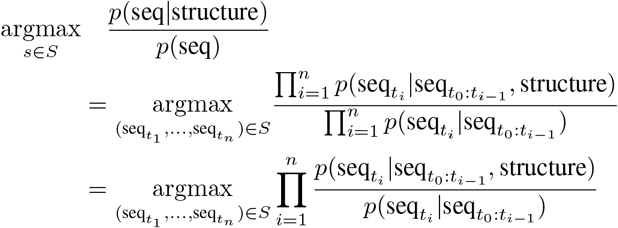

where 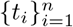 is a decoding order chosen by the user. We can also hold any position in the sequence fixed, where seq_*i*_ is assigned, the *i*-th term in the product passes out of the argmax operator, and the subsequent tokens in the decoding order are conditioned on seq_*i*_.

In NanoLuc experiments, residues essential to the protein function are fixed. In this case we decode using a “proximity” decode order in order to prioritize selection of residues that stabilize the active site. We decode using beam search (*n* = 128 beams).

In WW experiments, we decode with an “N to C” decode order, where residues are decoded in order from N-terminus to C-terminus. In this case we decode with greedy search.

#### 2.3.3. Numerical Trick

Because small probabilities in the denominator can result in large probability ratios, we add a small quantity *τ* (= 0.002) to the numerator and denominator probabilities, calculate the probability ratio, and re-normalize to sum to one. We find that this helps to avoid selecting residues with very low probability according to ProteinMPNN (see Figure S3).

## 3. Methods

As shown in the theory section, the Boltzmann probability objective is linked to protein stability and conformational specificity. An additional goal was to design soluble sequences, unlike previous approaches that used the Boltzmann probability objective. Thus, we evaluated designed sequences for stability, conformational specificity, and solubility.

We additionally validated designed sequences by comparing their AlphaFold-predicted structures to the AlphaFold-predicted structure of the wild type sequence (Jumper et al., 2021).

### 3.1. Evaluation of stability and solubility for NanoLuc

We validate the stability and solubility of sequences designed by the BayesDesign algorithm for the NanoLuc enzyme.

NanoLuc has been heavily engineered to increase its enzymatic activity. (Hall et al., 2012) mutated 16 residues close to the active site, resulting in an increase in enzymatic activity by several orders of magnitude. We redesign the enzyme while conserving various combinations of the active site and engineered residues, shown in Figure S5 and Table S1. We evaluate the change in stability and solubility relative to the wild type sequence.

In vitro validation experiments are conducted using cell-free protein synthesis (CFPS), a versatile expression system that is well-suited for experimental in vitro investigations of de novo designed proteins (Schinn et al., 2016) (Dopp et al., 2019) (Woodrow et al., 2006). CFPS is used in this work to obtain temperature-dependent protein solubility, which can be used as an indication of relative thermodynamic stability (Jarzab et al., 2020) (van Koningsveld et al., 2001) (Liu et al., 2015). The BayesDesign NanoLuc mutants and wild type NanoLuc are synthesized in vitro.

The synthesized mutants are heat treated at temperatures ranging from 37 to 70 degrees C. After heat treatment, the samples are centrifuged, and soluble protein in the supernatant is measured. Experimental methods are detailed in the supplementary information.

### 3.2. Evaluation of conformational specificity for WW

We also evaluate the conformational specificity of designed sequences by redesigning the WW domain of human protein Pin 1.

The structure of this short 34-residue peptide has been widely characterized (Lawrence et al., 2014) and it is known to follow a two-state model of unfolding. In this study we use the reversibility of WW as a proxy for conformational specificity. Sequences with high conformational specificity tend to refold to the native conformation after heat treatment (i.e. are reversible); others fail to recover the native state due to falling into alternate conformations upon refolding (Zhu et al., 2015). WW is chosen due to its poor reversibility, as low as 40% in some design studies (Xiao et al., 2019). We redesign the WW sequence with BayesDesign with the hypothesis that maximizing the Boltzmann probability objective will increase the conformational specificity of the native state.

We evaluate conformational specificity via the reversibility of the circular dichroism (CD) spectrum. CD measures the wavelength spectrum over a heating and cooling process. WW shows a characteristic peak at 227 nm that disappears upon thermal denaturation. We evaluate the reversibility of WW (and designed sequences) by measuring the percent of the starting CD signal at 227 nm recovered after equilibration at 95°C and cooling back to 25°C as in (Rago et al., 2015).

We compare the reversibility of the wild type sequence, a sequence designed by BayesDesign, and a sequence designed by ProteinMPNN. The rationale for using ProteinMPNN as a baseline is to compare the reversibility of sequences designed using the probability ratio of ProteinMPNN and XLNet (i.e. BayesDesign) versus sequences designed using the probabilities of ProteinMPNN alone.

## 4. Results

As preliminary validation, we evaluated AlphaFold-predicted structures for NanoLuc wild type and designed sequences. AlphaFold-predicted structures for designed sequences aligned closely with the predicted structure for the wild type sequence (see Figure S4).

In terms of stability, designed sequences showed increased stability relative to the original sequence, while maintaining solubility. We found increased stability in mutants M1-M4 relative to the original hand-engineered WT sequence (see Figure 3). While the hand-engineered WT showed increased stability relative to the original luciferase enzyme (Hall et al., 2012), our method afforded a further increase in stability. This shows that considering the full sequence when redesigning a protein offers opportunity for a significant increase in stability. Deep learning approaches like BayesDesign make it possible to quickly redesign the entire protein sequence to increase stability, while taking advantage of stability patterns learned from data. The observed stability increases are particularly notable in the context of a reported 2% success rate of random substitution mutants to increase stability (Pokala & Handel, 2005) (Foit et al., 2009) (Broom et al., 2012) (Araya et al., 2012) (Broom et al., 2017).

**Figure 3.**
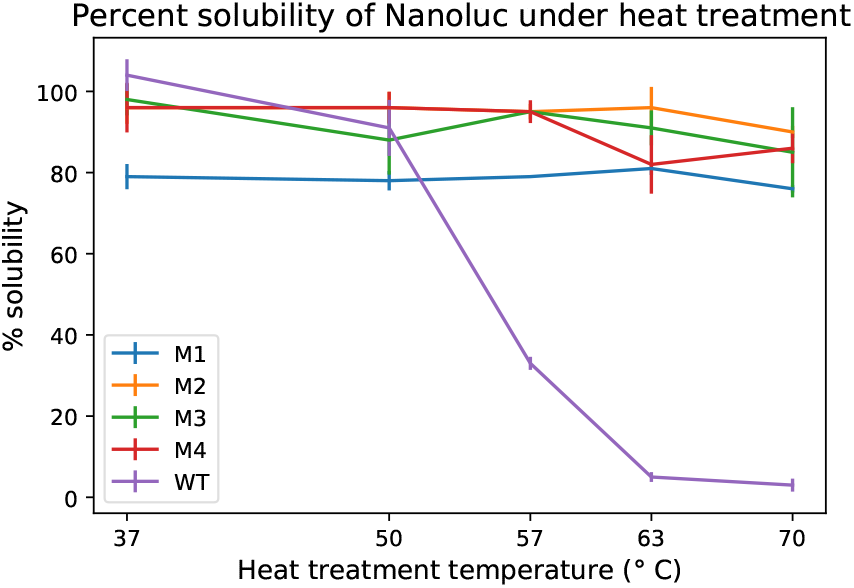
Designed NanoLuc mutants retained solubility at high temperatures, whereas the wild type denatured over heat treatment in the same range, indicating that BayesDesign increased the stability of the native structure in all mutants.

Sequence designs that improve stability often do so at the expense of solubility (Broom et al., 2017). Protein solubility is a desirable, if not essential, characteristic of active proteins for industrial, medicinal, and other applications (Broom et al., 2017) (Qing et al., 2022) (van Koningsveld et al., 2001). From a biochemical perspective, randomly generated protein sequences are rarely soluble (Wei et al., 2003) and random substitution mutations typically decrease protein solubility (Broom et al., 2017). From an optimization perspective, previous gradient-based approaches to optimizing the *p*(structure|seq) objective often resulted in insoluble sequences (Dauparas et al., 2022), requiring modification of the objective in order to obtain soluble sequence designs (Norn et al., 2021).

All designed sequences in this work (M1-M4) are soluble, ranging from 78 to 92 percent solubility (see Figure 4). To our knowledge, this work is the first to present soluble protein designs using an algorithm that optimizes the unmodified *p*(structure|seq) objective.

**Figure 4.**
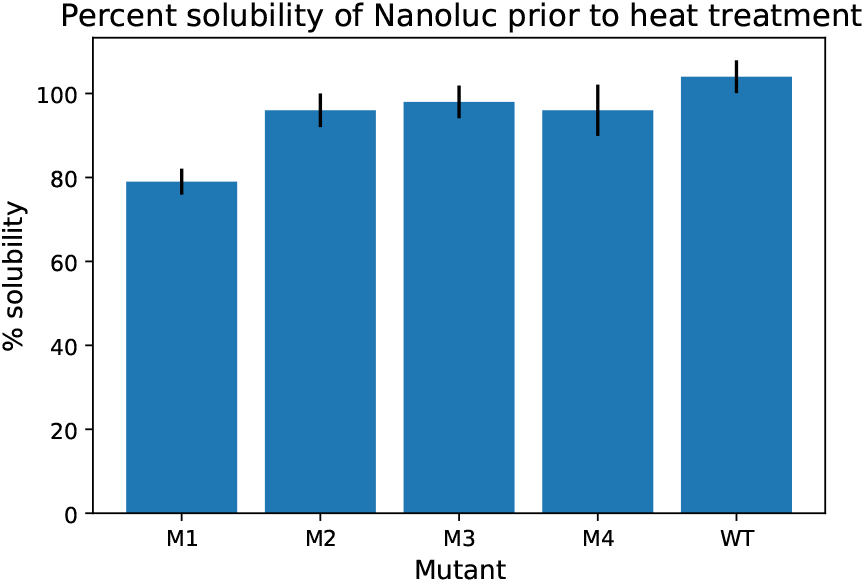
All NanoLuc mutants were found to be at least 75% soluble.

These stabilization results come with an important caveat: the BayesDesign-designed sequences removed almost all enzymatic activity of NanoLuc, as shown in Figure S7. This points to the potential pitfalls of redesigning proteins whose function is sensitive to changes in structure and flexibility. First, changing a large number of residues is more likely to change a residue essential to protein function. Second, the theory introduced in section 2.1.1 shows that the Boltzmann probability objective is linked to stability and conformational specificity, which may limit the flexibility needed to attain a catalytic conformation. The elimination of enzymatic activity indicates that the Boltzmann probability objective is not well suited for designing enzymes whose function depends on flexibility.

We also evaluated the BayesDesign algorithm on the short peptide WW. Solid-phase peptide synthesis was used to synthesize the native WW sequence, the BayesDesign redesign as well as a redesign by ProteinMPNN (see Tables S2 and S3 and Figure S10 for sequences and characterization).

AlphaFold validation of BayesDesign and ProteinMPNN-designed sequences showed close agreement between ProteinMPNN and the wild type structures and partial agreement between the BayesDesign structure and the wild type (see Figure 5).

**Figure 5.**
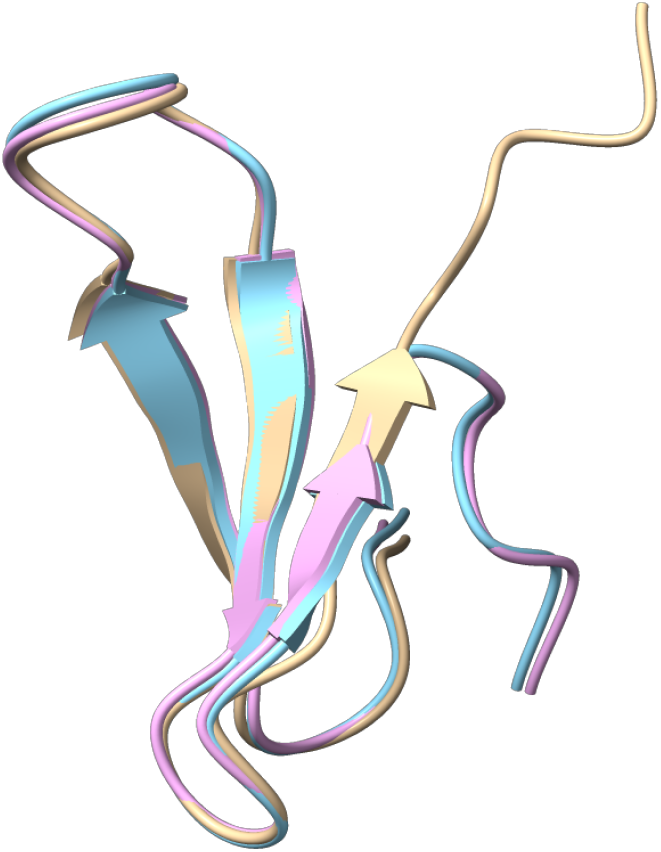
AlphaFold-predicted structures for the WW wild type (magenta), BayesDesign (tan), and ProteinMPNN (cyan) sequences.

However, a wavelength scan at 25°C revealed that neither ProteinMPNN nor BayesDesign sequences had the characteristic peak at 227 nm like that of WW (see Figure 6). The ProteinMPNN sequence had a peak at 230 nm, making it sufficiently different to be considered a different conformation. The BayesDesign sequence had no peak but rather a minimum at 203 nm. This indicates that both ProteinMPNN and BayesDesign failed to take on the exact structure of WW. This limits the effectiveness of the reversibility study, as only the 227 nm peak characteristic to native WW was tracked during the full heat treatment process for all peptides.

**Figure 6.**
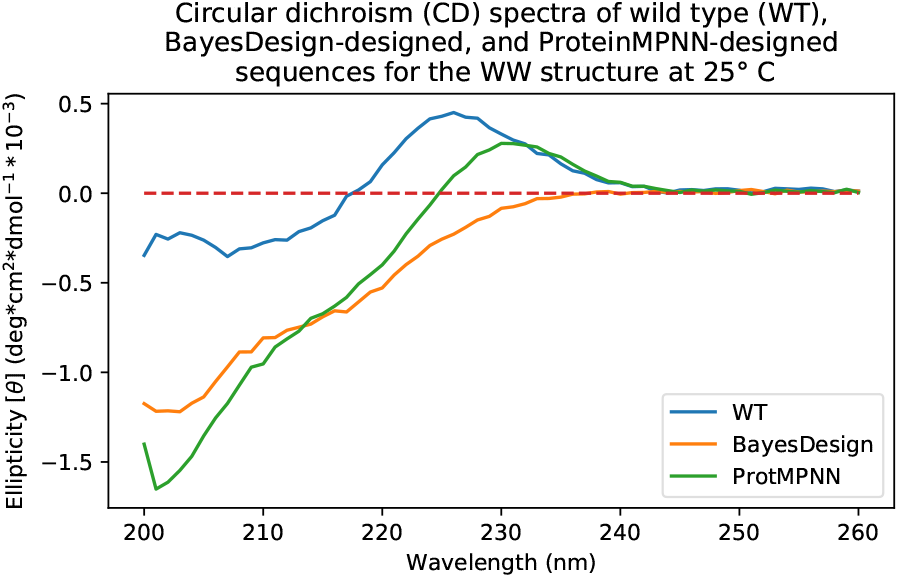
Circular dichroism (CD) profile of wild type (blue), BayesDesign (orange), and ProteinMPNN (green) sequences at 25°C. ProteinMPNN generally follows the ellipticity trends of the wild type structure, whereas BayesDesign does not have the same distinguishing peak at 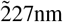.

Heating the BayesDesign peptide to 95 °C showed an ellipticity shift of the minimum at 203 nm, indicating some denaturation. After cooling, the minimum at 203 nm was completely recovered, showing that the BayesDesign peptide was 100% reversible for its adopted conformation at its minimum of 203 nm (see Figure S9) whereas native WW was only 56% reversibile at its 227 nm peak (see Figure S8). The high reversibility of the BayesDesign peptide indicates high conformational specificity, but not for the intended structure. A limited amount of peptide prevented an additional experiment to measure the reversibility of the ProteinMPNN peptide at its 230 nm peak.

## 5. Discussion

The results of this investigation were mixed, and offer some important lessons about pitfalls in using probabilistic models for protein sequence design.

### 5.1. Pitfall 1: Using the wrong objective

- If the goal is to increase protein stability, Boltzmann probability is a good objective but not as good as a probability model for stability.
- If the goal is to increase conformational specificity, Boltzmann probability is a good objective but not as good as a probability model for conformational specificity.
- If the goal is to increase enzymatic activity, Boltzmann probability may be the wrong objective because both stability and specificity limit the flexibility of enzymes, which is often key to their function.
- Maximizing the likelihood of the sequence *p*(seq |structure) is not guaranteed to increase stability or specificity, but may have other theoretical properties. We leave the investigation of the theoretical properties of this objective to future work.

### 5.2. Pitfall 2: Using the wrong probability model for the objective

If the model is trained on data, the training data may or may not represent the behavior you want to model. We believe that this is the main source of error in BayesDesign. ProteinMPNN was trained on static protein crystal structures in the PDB. ProtXLNet was trained on the UniRef100 database (Consortium, 2022). Both of these models capture the joint probability of protein sequences and structures as selected for by nature. Within that joint distribution, ProteinMPNN predicts sequence conditioned on structure, and ProtXLNet predicts sequence marginalized (summed) over all structures. ProteinMPNN treats protein structure as fixed, conditioning on a point estimate of protein structure as opposed to a sample from the Boltzmann distribution over protein structures.

As such, BayesDesign can be thought to model the probability distribution over fixed crystal structures for a given sequence, which is not necessarily the same as the Boltzmann distribution. We believe that this is why BayesDesign fails to design a sequence that folds to the native structure of WW.

### 5.3. Pitfall 3: Using a poor optimization scheme for the probability model

One positive lesson from this work is the optimization scheme. Using greedy and beam-search decoding resulted in soluble, foldable protein sequences. Gradient-based optimization schemes appear to be too good at finding minima, to the point where they find improbable sequences that highlight the failings of the forward prediction model. This could be viewed as a failure of the forward prediction model (given these poor sequences, the forward prediction model should give low probability for the desired structure), but the optimization algorithm is related to the choice of model. In this work, the combined choice of model and optimization scheme was found to be an effective choice for designing the soluble sequences.

## 6. Conclusion

We introduce a probabilistic framework for predicting the effects of the Boltzmann probability objective function *p*(structure|seq) on protein stability and conformational specificity. We introduce a BayesDesign, a new protein design algorithm to maximize this design objective by applying Bayes’ rule to unconditional and structure-conditioned autoregressive sequence models.

Compared to other methods that maximize *p*(structure| seq), our method seems to suffer less from adversarial sequence problems in the optimization procedure. Compared to methods that maximize *p*(seq| structure), we have theoretical guarantees ensuring that designed sequences should increase protein stability and/or conformational specificity.

We show that redesigning the NanoLuc enzyme with this objective increases its stability. For the WW peptide, the redesigned sequence fails to fold to the correct structure, and we identify several possible model improvements to address in future work.

## References

Alford, R. F., Leaver-Fay, A., Jeliazkov, J. R., O’Meara, M. J., DiMaio, F. P., Park, H., Shapovalov, M. V., Renfrew, P. D., Mulligan, V. K., Kappel, K., Labonte, J. W., Pacella, M. S., Bonneau, R., Bradley, P., Dunbrack, R. L., Das, R., Baker, D., Kuhlman, B., Kortemme, T., and Gray, J. J. The rosetta all-atom energy function for macromolecular modeling and design. Journal of Chemical Theory and Computation, 13(6):3031–3048, Jun 2017. ISSN 1549-9618. doi: 10.1021/acs.jctc.7b00125. URL https://doi.org/10.1021/acs.jctc.7b00125.

Anand, N., Eguchi, R., Mathews, I. I., Perez, C. P., Derry, A., Altman, R. B., and Huang, P.-S. Protein sequence design with a learned potential. Nature Communications, 13(1):746, Feb 2022. ISSN 2041-1723. doi: 10.1038/s41467-022-28313-9. URL https://doi.org/10.1038/s41467-022-28313-9.

Anishchenko, I., Pellock, S. J., Chidyausiku, T. M., Ramelot, T. A., Ovchinnikov, S., Hao, J., Bafna, K., Norn, C., Kang, A., Bera, A. K., DiMaio, F., Carter, L., Chow, C. M., Montelione, G. T., and Baker, D. De novo protein design by deep network hallucination. Nature, 600(7889): 547–552, Dec 2021. ISSN 1476-4687. doi: 10.1038/s41586-021-04184-w. URL https://doi.org/10.1038/s41586-021-04184-w.

Araya, C. L., Fowler, D. M., Chen, W., Muniez, I., Kelly, J. W., and Fields, S. A fundamental protein property, thermodynamic stability, revealed solely from large-scale measurements of protein function. Proceedings of the National Academy of Sciences, 109(42):16858–16863, 2012. doi: 10.1073/pnas.1209751109. URL https://www.pnas.org/doi/abs/10.1073/pnas.1209751109.

Broom, A., Doxey, A. C., Lobsanov, Y. D., Berthin, L. G., Rose, D. R., Howell, P. L., McConkey, B. J., and Meiering, E. M. Modular evolution and the origins of symmetry: Reconstruction of a three-fold symmetric globular protein. Structure, 20(1):161–171, Jan 2012. ISSN 0969-2126. doi: 10.1016/j.str.2011.10.021. URL https://doi.org/10.1016/j.str.2011.10.021.

Broom, A., Jacobi, Z., Trainor, K., and Meiering, E. M. Computational tools help improve protein stability but with a solubility tradeoff. Journal of Biological Chemistry, 292(35):14349–14361, Sep 2017. ISSN 0021-9258. doi: 10.1074/jbc.M117.784165. URL https://doi.org/10.1074/jbc.M117.784165.

Consortium, T. U. UniProt: the Universal Protein Knowledgebase in 2023. Nucleic Acids Research, 11 2022. ISSN 0305-1048. doi: 10.1093/nar/gkac1052. URL https://doi.org/10.1093/nar/gkac1052.gkac1052.

Dahiyat, B. I. and Mayo, S. L. De novo protein design: Fully automated sequence selection. Science, 278(5335):82–87, 1997. doi: 10.1126/science.278.5335.82. URL https://www.science.org/doi/abs/10.1126/science.278.5335.82.

Dauparas, J., Anishchenko, I., Bennett, N., Bai, H., Ragotte, R. J., Milles, L. F., Wicky, B. I. M., Courbet, A., de Haas, R. J., Bethel, N., Leung, P. J. Y., Huddy, T. F., Pellock, S., Tischer, D., Chan, F., Koepnick, B., Nguyen, H., Kang, A., Sankaran, B., Bera, A. K., King, N. P., and Baker, D. Robust deep learning based protein sequence design using proteinmpnn. bioRxiv, 2022. doi: 10.1101/2022.06.03.494563. URL https://www.biorxiv.org/content/early/2022/06/04/2022.06.03.494563.

Dopp, J. L., Rothstein, S. M., Mansell, T. J., and Reuel, N. F. Rapid prototyping of proteins: Mail order gene fragments to assayable proteins within 24hours. Biotechnology and Bioengineering, 116 (3):667–676, 2019. doi: https://doi.org/10.1002/bit.26912. URL https://onlinelibrary.wiley.com/doi/abs/10.1002/bit.26912.

Elnaggar, A., Heinzinger, M., Dallago, C., Rihawi, G., Wang, Y., Jones, L., Gibbs, T., Feher, T., Angerer, C., Steinegger, M., Bhowmik, D., and Rost, B. Prottrans: Towards cracking the language of life’s code through selfsupervised deep learning and high performance computing, 2020. URL https://arxiv.org/abs/2007.06225.

Foit, L., Morgan, G. J., Kern, M. J., Steimer, L. R., von Hacht, A. A., Titchmarsh, J., Warriner, S. L., Radford, S. E., and Bardwell, J. C. Optimizing protein stability in vivo. Molecular Cell, 36(5):861–871, Dec 2009. ISSN 1097-2765. doi: 10.1016/j.molcel.2009.11.022. URL https://doi.org/10.1016/j.molcel.2009.11.022.

Goodfellow, I. J., Shlens, J., and Szegedy, C. Explaining and harnessing adversarial examples, 2014. URL https://arxiv.org/abs/1412.6572.

Hall, M. P., Unch, J., Binkowski, B. F., Valley, M. P., Butler, B. L., Wood, M. G., Otto, P., Zimmerman, K., Vidugiris, G., Machleidt, T., Robers, M. B., Benink, H. A., Eggers, C. T., Slater, M. R., Meisenheimer, P. L., Klaubert, D. H., Fan, F., Encell, L. P., and Wood, K. V. Engineered luciferase reporter from a deep sea shrimp utilizing a novel imidazopyrazinone substrate. ACS Chemical Biology, 7(11):1848–1857, Nov 2012. ISSN 1554-8929. doi: 10.1021/cb3002478. URL https://doi.org/10.1021/cb3002478.

Ingraham, J., Garg, V. K., Barzilay, R., and Jaakkola, T. Generative Models for Graph-Based Protein Design. Curran Associates Inc., Red Hook, NY, USA, 2019.

Ingraham, J., Baranov, M., Costello, Z., Frappier, V., Ismail, A., Tie, S., Wang, W., Xue, V., Obermeyer, F., Beam, A., and Grigoryan, G. Illuminating protein space with a programmable generative model. bioRxiv, 2022. doi: 10.1101/2022.12.01.518682. URL https://www.biorxiv.org/content/early/2022/12/02/2022.12.01.518682.

Jarzab, A., Kurzawa, N., Hopf, T., Moerch, M., Zecha, J., Leijten, N., Bian, Y., Musiol, E., Maschberger, M., Stoehr, G., Becher, I., Daly, C., Samaras, P., Mergner, J., Spanier, B., Angelov, A., Werner, T., Bantscheff, M., Wilhelm, M., Klingenspor, M., Lemeer, S., Liebl, W., Hahne, H., Savitski, M. M., and Kuster, B. Meltome atlas—thermal proteome stability across the tree of life. Nature Methods, 17(5):495–503, May 2020. ISSN 1548-7105. doi: 10.1038/s41592-020-0801-4. URL https://doi.org/10.1038/s41592-020-0801-4.

Jones, D. T. De novo protein design using pairwise potentials and a genetic algorithm. Protein Science, 3(4):567–574, 1994. doi: https://doi.org/10.1002/pro.5560030405. URL https://onlinelibrary.wiley.com/doi/abs/10.1002/pro.5560030405.

Jumper, J., Evans, R., Pritzel, A., Green, T., Figurnov, M., Ronneberger, O., Tunyasuvunakool, K., Bates, R., Žídek, A., Potapenko, A., Bridgland, A., Meyer, C., Kohl, S. A. A., Ballard, A. J., Cowie, A., Romera-Paredes, B., Nikolov, S., Jain, R., Adler, J., Back, T., Petersen, S., Reiman, D., Clancy, E., Zielinski, M., Steinegger, M., Pacholska, M., Berghammer, T., Bodenstein, S., Silver, D., Vinyals, O., Senior, A. W., Kavukcuoglu, K., Kohli, P., and Hassabis, D. Highly accurate protein structure prediction with alphafold. Nature, 596 (7873):583–589, Aug 2021. ISSN 1476-4687. doi: 10.1038/s41586-021-03819-2. URL https://doi.org/10.1038/s41586-021-03819-2.

Kuhlman, B., Dantas, G., Ireton, G. C., Varani, G., Stoddard, B. L., and Baker, D. Design of a novel globular protein fold with atomic-level accuracy. Science, 302(5649):1364–1368, 2003. doi: 10.1126/science.1089427. URL https://www.science.org/doi/abs/10.1126/science.1089427.

Lawrence, P. B., Gavrilov, Y., Matthews, S. S., Langlois, M. I., Shental-Bechor, D., Greenblatt, H. M., Pandey, B. K., Smith, M. S., Paxman, R., Torgerson, C. D., Merrell, J. P., Ritz, C. C., Prigozhin, M. B., Levy, Y., and Price, J. L. Criteria for selecting pegylation sites on proteins for higher thermodynamic and proteolytic stability. Journal of the American Chemical Society, 136(50):17547–17560, 2014. doi: 10.1021/ja5095183. URL https://doi.org/10.1021/ja5095183. PMID: 25409346.

Liu, J. L., Goldman, E. R., Zabetakis, D., Walper, S. A., Turner, K. B., Shriver-Lake, L. C., and Anderson, G. P. Enhanced production of a single domain antibody with an engineered stabilizing extra disulfide bond. Microbial Cell Factories, 14(1):158, Oct 2015. ISSN 1475-2859. doi: 10.1186/s12934-015-0340-3. URL https://doi.org/10.1186/s12934-015-0340-3.

Liu, Y. and Kuhlman, B. RosettaDesign server for protein design. Nucleic Acids Research, 34(suppl2) : W 235 – –W 238, 072006.ISSN 0305–1048.doi :. URL https://doi.org/10.1093/nar/gkl163.

Marshall, S. A. and Mayo, S. L. Achieving stability and conformational specificity in designed proteins via binary patterning. Journal of Molecular Biology, 305(3):619–631, 2001. ISSN 0022-2836. https://doi.org/11.1006/jmbi.2000.4319. URL https://www.sciencedirect.com/science/article/pii/S0022283600943195.

Norn, C., Wicky, B. I. M., Juergens, D., Liu, S., Kim, D., Tischer, D., Koepnick, B., Anishchenko, I., null null, Baker, D., and Ovchinnikov, S. Protein sequence design by conformational landscape optimization. Proceedings of the National Academy of Sciences, 118(11):e2017228118, 2021. 10.1073/pnas.2017228118. URL https://www.pnas.org/doi/abs/10.1073/pnas.2017228118.

Pokala, N. and Handel, T. M. Energy functions for protein design: Adjustment with protein–protein complex affinities, models for the unfolded state, and negative design of solubility and specificity. Journal of Molecular Biology, 347(1):203–227, 2005. ISSN 0022-2836. https://doi.org/10.1016/j.jmb.2004.12.019. URL https://www.sciencedirect.com/science/article/pii/S002228360401589X.

Qing, R., Hao, S., Smorodina, E., Jin, D., Zalevsky, A., and Zhang, S. Protein design: From the aspect of water solubility and stability. Chemical Reviews, 122 (18):14085–14179, 2022. 10.1021/acs.chemrev.1c00757. URL https://doi.org/10.1021/acs.chemrev.1c00757. PMID: 35921495.

Rago, F., Saltzberg, D., Allen, K. N., and Tolan, D. R. Enzyme substrate specificity conferred by distinct conformational pathways. Journal of the American Chemical Society, 137(43):13876–13886, 2015. 10.1021/jacs.5b08149. URL https://doi.org/10.1021/jacs.5b08149. PMID: 26440863.

Roney, J. P. and Ovchinnikov, S. State-of-the-art estimation of protein model accuracy using alphafold. Phys. Rev. Lett., 129:238101, Nov 2022. 10.1103/Phys-RevLett.129.238101. URL https://link.aps.org/doi/10.1103/PhysRevLett.129.238101.

Schinn, S.-M., Broadbent, A., Bradley, W. T., and Bundy, C. Protein synthesis directly from pcr: progress and applications of cell-free protein synthesis with linear dna. New Biotechnology, 33(4):480–487, 2016. ISSN 1871-6784. https://doi.org/10.1016/j.nbt.2016.04.002. URL https://www.sciencedirect.com/science/article/pii/S187167841630005X.

Simons, K. T., Kooperberg, C., Huang, E., and Baker, D. Assembly of protein tertiary structures from fragments with similar local sequences using simulated annealing and bayesian scoring functions11edited by f. e. cohen. Journal of Molecular Biology, 268(1):209–225, 1997. ISSN 0022-2836. https://doi.org/10.1006/jmbi.1997.0959. URL https://www.sciencedirect.com/science/article/pii/S0022283697909591.

van Koningsveld, G. A., Gruppen, H., de Jongh, H. H. J., Wijngaards, G., van Boekel, M. A. J. S., Walstra, P., and Voragen, A. G. J. Effects of ph and heat treatments on the structure and solubility of potato proteins in different preparations. Journal of Agricultural and Food Chemistry, 49(10):4889–4897, 2001. 10.1021/jf010340j. URL https://doi.org/10.1021/jf010340j. PMID: 11600040.

Wang, J., Lisanza, S., Juergens, D., Tischer, D., Watson, J. L., Castro, K. M., Ragotte, R., Saragovi, A., Milles, L. F., Baek, M., Anishchenko, I., Yang, W., Hicks, D. R., Expòsit, M., Schlichthaerle, T., Chun, J.-H., Dauparas, J., Bennett, N., Wicky, B. I. M., Muenks, A., DiMaio, F., Correia, B., Ovchinnikov, S., and Baker, D. Scaffolding protein functional sites using deep learning. Science, 377(6604):387–394, 2022. 10.1126/science.abn2100. URL https://www.science.org/doi/abs/10.1126/science.abn2100.

Watson, J. L., Juergens, D., Bennett, N. R., Trippe, B. L., Yim, J., Eisenach, H. E., Ahern, W., Borst, A. J., Ragotte, R. J., Milles, L. F., Wicky, B. I. M., Hanikel, N., Pellock, S. J., Courbet, A., Sheffler, W., Wang, J., Venkatesh, P., Sappington, I., Torres, S. V., Lauko, A., De Bortoli, V., Mathieu, E., Barzilay, R., Jaakkola, T. S., DiMaio, F., Baek, M., and Baker, D. Broadly applicable and accurate protein design by integrating structure prediction networks and diffusion generative models. bioRxiv, 2022. 10.1101/2022.12.09.519842. URL https://www.biorxiv.org/content/early/2022/12/10/2022.12.09.519842.

Wei, Y., Kim, S., Fela, D., Baum, J., and Hecht, M. H. Solution structure of a ¡i¿de novo¡/i¿ protein from a designed combinatorial library. Proceedings of the National Academy of Sciences, 100(23):13270–13273, 2003. 10.1073/pnas.1835644100. URL https://www.pnas.org/doi/abs/10.1073/pnas.1835644100.

Woodrow, K. A., Airen, I. O., and Swartz, J. R. Rapid expression of functional genomic libraries. Journal of Proteome Research, 5(12):3288–3300, 2006. 10.1021/pr050459y. URL https://doi.org/10.1021/pr050459y. PMID: 17137330.

Xiao, Q., Draper, S. R. E., Smith, M. S., Brown, N., Pugmire, N. A. B., Ashton, D. S., Carter, A. J., Lawrence, E. E. K., and Price, J. L. Influence of pegylation on the strength of protein surface salt bridges. ACS Chemical Biology, 14(7):1652–1659, 2019. 10.1021/acschem-bio.9b00432. URL https://doi.org/10.1021/acschembio.9b00432. PMID: 31188563.

Yang, J., Anishchenko, I., Park, H., Peng, Z., Ovchinnikov, S., and Baker, D. Improved protein structure prediction using predicted interresidue orientations. Proceedings of the National Academy of Sciences, 117(3):1496–1503, 2020. 10.1073/pnas.1914677117. URL https://www.pnas.org/doi/abs/10.1073/pnas.1914677117.

Zhu, G.-F., Ren, S.-Y., Xi, L., Du, L.-F., and Zhu, X.-F. Temperature induced structural transitions from native to unfolded aggregated states of tobacco etch virus protease. Journal of Molecular Structure, 1082:80–90, 2015. ISSN 0022-2860. https://doi.org/10.1016/j.molstruc.2014.11.010. URL https://www.sciencedirect.com/science/article/pii/S0022286014011168.

