## Supplementary Information for "A probabilistic view of protein stability, conformational specificity, and design"

December 28, 2022

#### A Supplemental figures

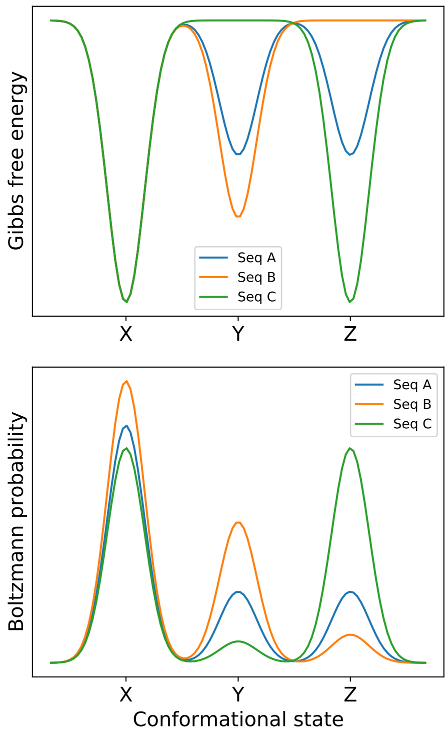


Figure 1: A case (i.e. a set of energy landscapes) where optimizing Boltzmann probability would result in the maximum stability sequence.

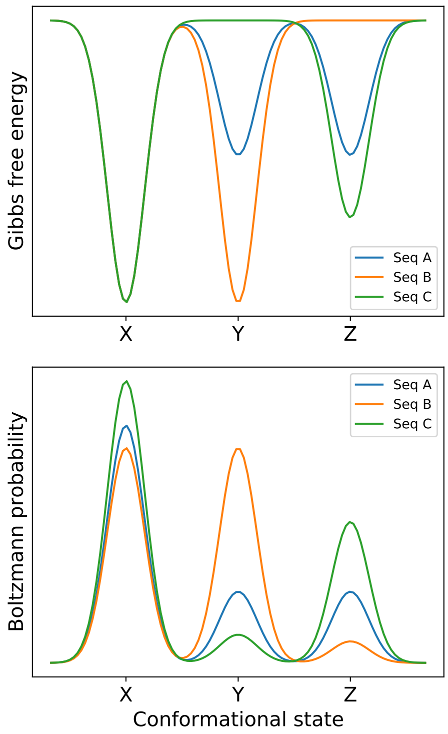


Figure 2: A case (i.e. a set of energy landscapes) where optimizing Boltzmann probability would result in the maximum specificity sequence.

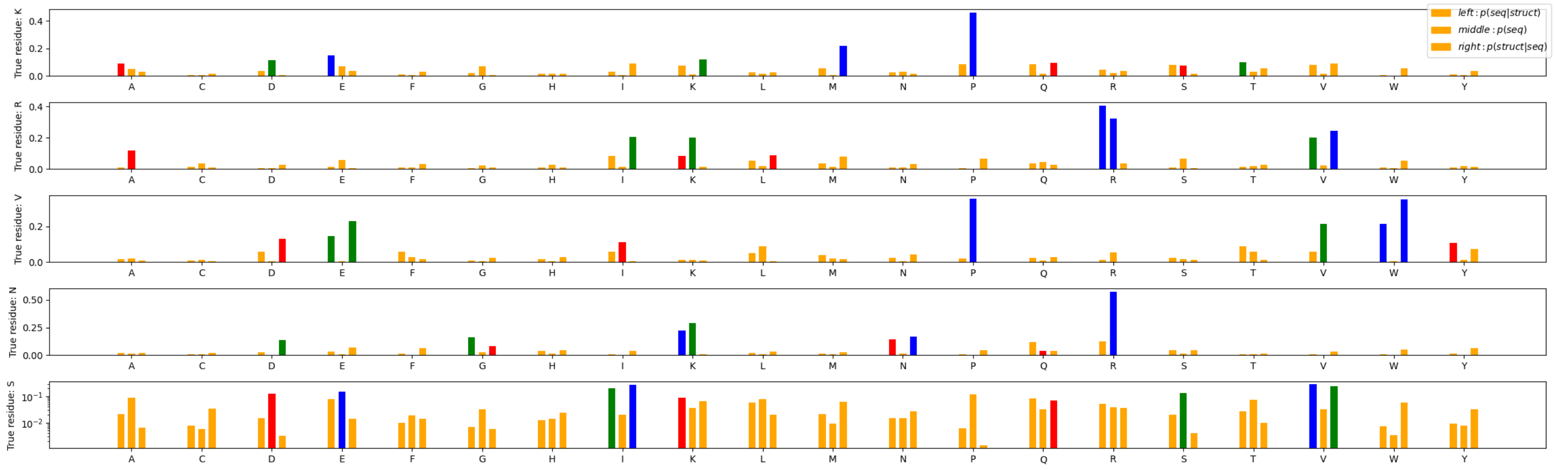


Figure 3: A visualization of probability shift induced by applying Bayes' rule to ProteinMPNN probabilities. Each row represents an evenly spaced residue in the WW peptide. Each column represents an amino acid. The first bar in each cluster corresponds to $p(\text{seq}|\text{structure})$; the second, to $p(\text{seq})$, and the third to $p(\text{seq}|\text{structure})/p(\text{seq})$, normalized across amino acids. The top score for each model is in blue, the second highest is in red, and the third highest is in green. In the second row, ProteinMPNN's best guess is arginine, but XLNet is in agreement, so the third model ends up choosing valine, which was originally the third choice of ProteinMPNN.

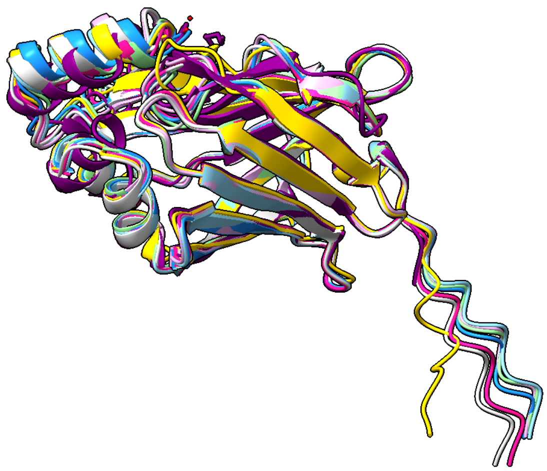

Figure 4: Structure alignment of the Alphafold-predicted structures of all mutants with the WT predicted structure.

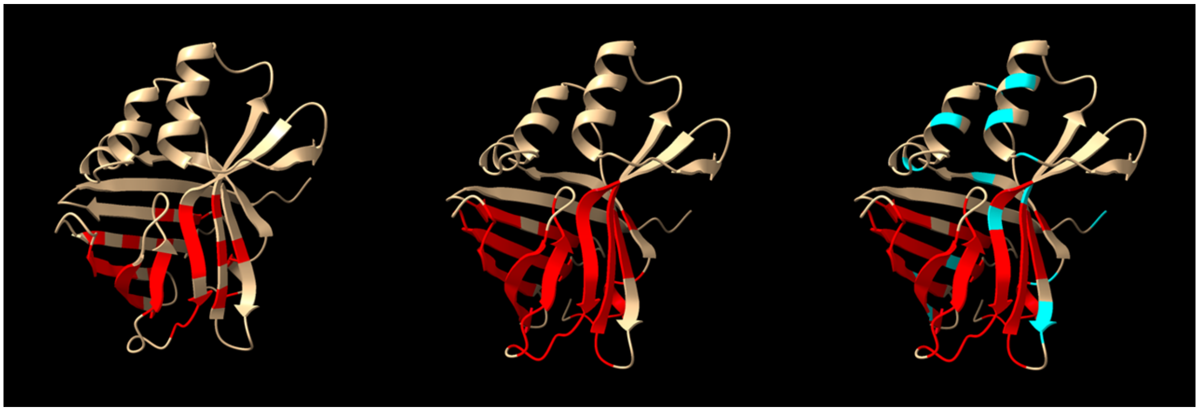

Figure 5: Several regions of the NanoLuc enzyme were considered in enzyme redesign. On the left, red highlights the residues around the enzyme active site. In the middle, the (red) region around the active site is expanded to include more residues. On the right, 14 previously hand-engineered residues are also highlighted in cyan. For our four mutants, we applied four design variations: 1) holding only the small active site amino acids fixed and designing all others (M1), 2) holding the the small active site amino acids and the engineered residues fixed and designing all others (M2), 3) holding only the large active site amino acids fixed and designing all others (M3), and 4) holding the the large active site amino acids and the engineered residues fixed and designing all others (M4).

```
PDB_5IBO     -----SDNMVFTLEDFVGDWRQTAGYNLDQVLEQGGVSSLFQNLGVSVTPIQRIVLSGEN    55
Wild_Type    MWSHPQFEKVFTLEDFVGDWRQTAGYNLDQVLEQGGVSSLFQNLGVSVTPIQRIVLSGEN    60
                   . : ***************************************************

PDB_5IBO     GLKIDIHVIIPYEGLSGDQMGQIEKIFKVVYPVDDHHFKVILHYGTLVIDGVTPNMIDYF    115
Wild_Type    GLKIDIHVIIPYEGLSGDQMGQIEKIFKVVYPVDDHHFKVILHYGTLVIDGVTPNMIDYF    120
             ************************************************************

PDB_5IBO     GRPYEGIAVFDGKKITVTGTLWNGNKIIDERLINPDGSLLFRVTINGVTGWRLCERILA     174
Wild_Type    GRPYEGIAVFDGKKITVTGTLWNGNKIIDERLINPDGSLLFRVTINGVTGWRLCERILA     179
             ***********************************************************
```

Figure 6: Comparison of the NanoLuc sequence referred to as wild type (WT) in this paper to the sequence of PDB 5IBO.

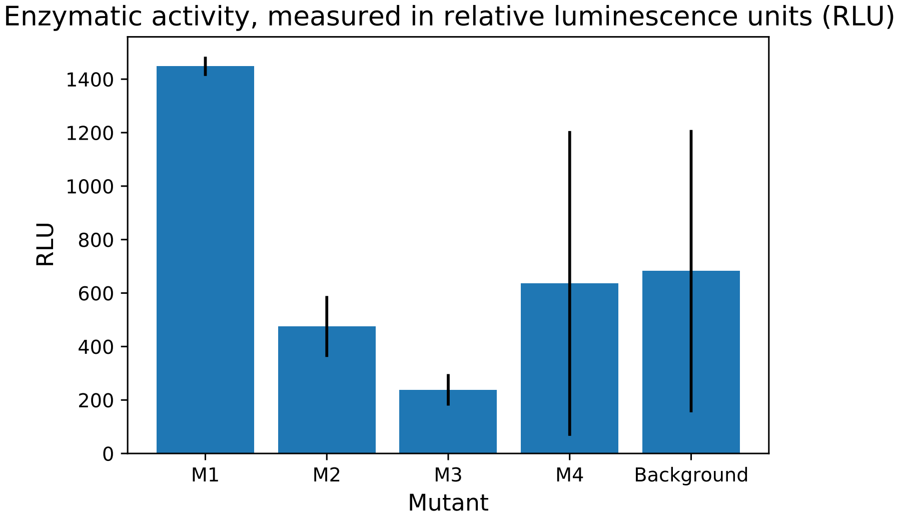


Figure 7: NanoLuc activity, measured in relative luminescence units (RLU). In the process of stabilizing NanoLuc, enzymatic activity was essentially eliminated. M1 has non-zero activity, but for comparison, the WT enzyme has activity of 158277 RLU, so an increase of 100% above background reaction activity is comparatively negligible.

Table 1: Wild type and designed sequences for NanoLuc. Masked positions are designed.

| | |
|---|---|
| WT | |
| Sequence | MWSHPQFEKVFTLEDFVGDWRQTAGYNLDQVLEQGGVSSLFQNLGVSVTPIQRIVLSG<br>ENGLKIDIHVIIPYEGLSGDQMGQIEKIFKVVYPVDDHHFKVILHYGTLVIDGVTPNM<br>IDYFGRPYEGIAVFDGKKITVTGTLWNGNKIIDERLINPDGSLLFRVTINGVTGWRLC<br>ERILA |
| BayesDesign - M1 | |
| Masked sequence | MWSHPQFEKV--L-------------------------------------I-------<br>E-G-K------------------------------------VIL-Y-T-----V-P--<br>----GRPYE-I----G-KITV------G-K-I-------D-S-L--------------<br>----- |
| Designed sequence | MWSHPQFEKVLTLDDFVGNWRMVSQWNIPAVLREMGMPPFLIDLWCATTPIWVITKYG<br>ENGLKVDVHMVIPKEGLTPEQMRYLQAMFGHMTQVDETHFQVILDYGVFIINGTSKNC<br>KDFMNRPFEVNTTFDGKKLTMTGTLWNGKKFVMTFEILPDGHLRYTVDVNGVKGWMIL<br>ERVEP |
| BayesDesign - M2 | |
| Masked sequence | MWSHPQFEKV--LE------R------L--------V-----N-------I-RI----<br>E-G-KI-------------D---Q--K--------------VIL-Y-T-----V-P--<br>----GRPYE-I----G-KITV------G-K-I-------D-S-L--------------<br>-R--- |
| Designed sequence | MWSHPQFEKVLKLEDFVGDWRRVDSWNLPEVLKAMGVPQFFINLFCQTQPIWRISKHG<br>EKGLKIQMIMRIPKQGLTPDQMAQIQKTFKHVQDIDDQHFQVILDYGTLIIDGVSPNC<br>KDFLGRPYEGICKFDGKKITVTGTLPNGNKFIWTMEILDDGSLLFTVDVNGVKGYMIL<br>ERVEP |
| BayesDesign - M3 | |
| Masked sequence | MWSHPQFEKVFTLE--V-------------------------------TPI-----GE<br>NGLKI---------------------------------FKVILHYGTLV-DGVTPNM-<br>--FGRPYEGI----GKKITVTG--WNGNKIIDE-----DGS-LFR-------------<br>---- |
| Designed sequence | MWSHPQFEKVFTLEDFVGDWRLVSKQNMAAVLREMGAPDFLIQLYLQCTPIFHITKSG<br>ENGLKIDVEMIIPKAGLTPEQMCYLQKMFKHMEPVDENHFKVILHYGTLVIDGVTPNM<br>KDAFGRPYEGICKFDGKKITVTGTLWNGNKIIDEYEILPDGSLLFRRTVNGVTGWMKL<br>ERVEP |
| BayesDesign - M4 | |
| Masked sequence | MWSHPQFEKVFTLE--V---R------L--------V-----N-----TPI-RI--SG<br>ENGLKI-------------D---Q--K------------FKVILHYGTLV-DGVTPNM<br>---FGRPYEGI----GKKITVTG--WNGNKIIDE-----DGS-LFR------------<br>-R--- |
| Designed sequence | MWSHPQFEKVFTLEDFVGDWREVDRWNLADVLKAMGVPQFLINLYMSCTPIWRITKSG<br>ENGLKIDVEMIIPKQGLTEDQLQQIKKIFQHVEDVDDNHFKVILHYGTLVIDGVTPNM<br>KDWFGRPYEGICKFDGKKITVTGTLWNGNKIIDEFEILPDGSLLFRTTVNGVTGYRIL<br>ERVEP |

Table 2: Wild type and designed sequences for WW. Masked positions are designed.

| | |
|---|---|
| WT | |
| Sequence | KLPPGWEKRMSRSSGRVYYFNHITNASQFERPSG |
| ProteinMPNN | |
| Masked sequence | ---------------------------------G |
| Designed sequence | TLPEGWVEVVDPETGEKKYYNTKTKEVTSEKPVG |
| BayesDesign | |
| Masked sequence | ---------------------------------G |
| Designed sequence | TLPEHWVKRKDPKTGQWIYENTKTHETLAQKWQG |

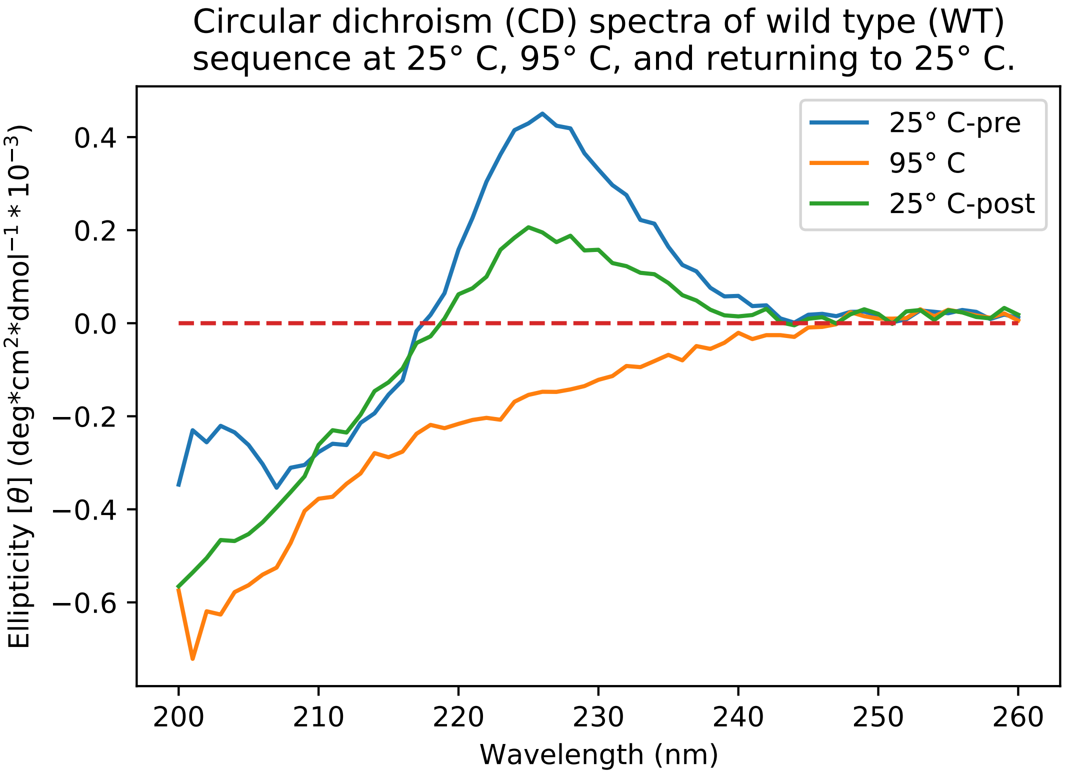


Figure 8: WW has a characteristic peak at $\tilde{2}27$nm, and has low reversibility of 56% - it fails to recover its original CD spectrum after heat treatment and cooling.

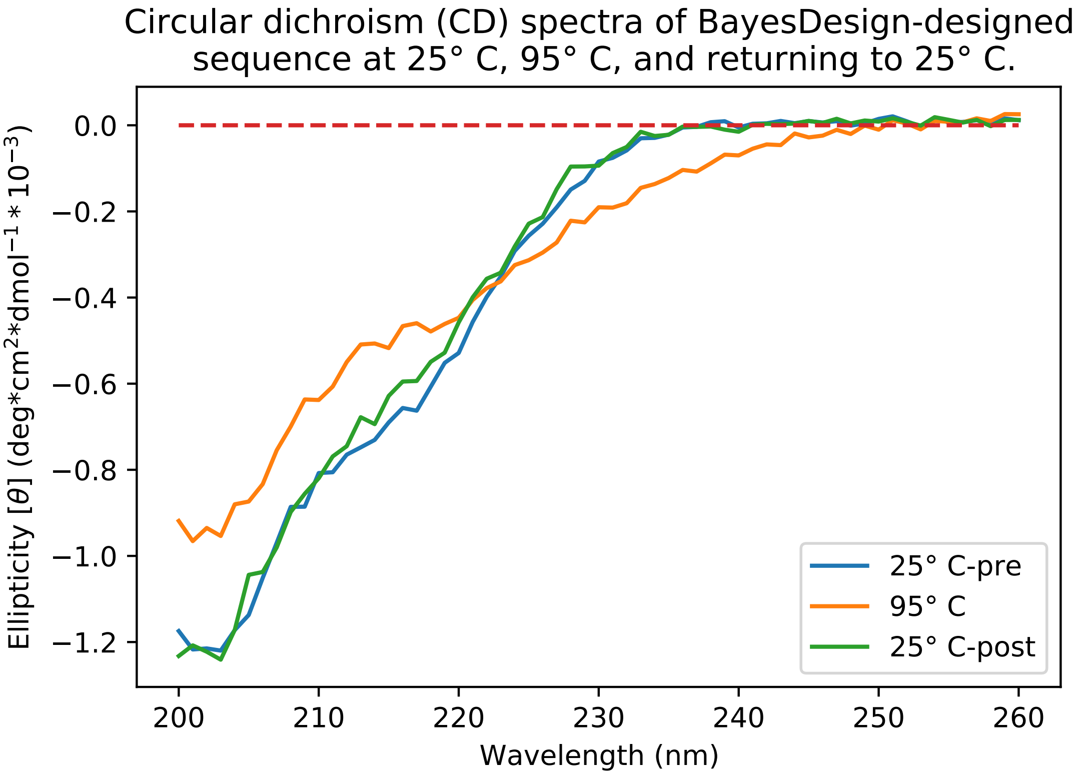


Figure 9: Although the BayesDesign design for WW does not follow the WT CD profile exactly, its profile shifts when heated and regains its original profile when returned to 25° C. This suggests that the designed sequence adopts a folded conformation (not the same conformation as the WT) with high conformational specificity.

Table 3: Mass spectrum data for synthesized WW mutants

| Peptide Name | m/z | Expected Mass | Observed Mass |
|---|---|---|---|
| Wild type | 4 | 996.2513 | 996.2494 |
| ProteinMPNN | 3 | 1279.9868 | 1279.9804 |
| BayesDesign | 3 | 1378.7160 | 1378.7075 |

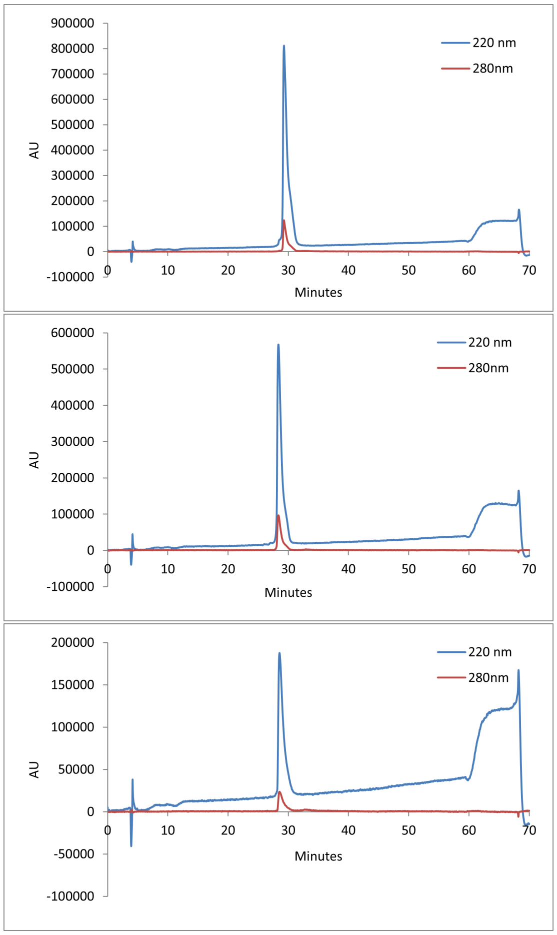


Figure 10: Analytical HPLC trace for WW wild type (top), BayesDesign (middle), and ProteinMPNN (bottom).

### B Experimental methods

#### B.1 NanoLuc synthesis and assays

##### B.1.1 DNA design

The NanoLuc protein sequence referred to as wild type in this work is the NanoLuc (PDB: 5IBO) amino acid sequence with an added N-terminal Strep tag as shown in Supplemental Figure 6. DNA sequences to express wild type NanoLuc and the BayesDesign mutants were designed to include the following regions: Forward primer binding site, non-transcribed spacer, T7 promoter, Ribosome Binding Site, protein-coding region, T7 terminator, non-transcribed spacer, reverse primer binding site. The DNA protein-coding region for each mutant and the wild type control were generated from the amino acid sequences of each mutant and the wild type control. The DNA codon choice was optimized for E coli (ThermoFisher GeneArt) and DNA gene fragments were constructed by Twist Bioscience (San Francisco, CA) and amplified by Q5 PCR (NEB, Ipswich, MA, USA).

##### B.1.2 Cell-free protein synthesis

Cell-free protein synthesis (CFPS) was conducted as described in [Hunt et al., 2022]. Cell extract was prepared using BL21-Star™ (DE3) E coli cells (Invitrogen, Carlsbad, CA) cultured in 2xYT media. The cells were induced with isopropyl $\beta$-d-1-thiogalactopyranoside (IPTG) at OD600 0.5 to 0.7, harvested at OD600 of 2 to 4, washed, and lysed with 3 passes through an Avestin Emulsiflex B-15 homogenizer (Avestin, Ottawa, Canada) at 21000 psi. Lysate was centrifuged at 12000 RCF for 30 minutes and the supernatant was harvested as cell extract, which is used at 25% (v/v) in CFPS reactions. For DNA template in CFPS reactions, unpurified PCR product was added at 33% (v/v). A PANOx-SP mixture of nucleotides, amino acids, energy substrates and other small molecules was added using reaction concentrations described in [Jewett and Swartz, 2004]. For yield determination, 2 $\mu$L samples of unpurified CFPS reaction product were dried on filter paper, precipitated and washed with TCA, and measured with a scintillation counter. Yields were calculated based on the percentage of incorporated 14C-leucine, which was present at a concentration of 5$\mu$M in the initial reaction preparation [Bundy and Swartz, 2010].

##### B.1.3 Identification of active site

Using Amber's ff14SB force field and OpenMM's molecular dynamics engine, we ran energy minimization on the NanoLuciferase crystal structure (5IBO). Docking runs were then performed with Vina, LeDock, and Plants 1.2, using the minimized protein and its ligand, furimazine. The area explored by the docking software consisted of a box that encompassed the entire beta barrel structure of the enzyme, based on the description of the active site in [Tomabechi et al., 2016]. Three MD simulations were then performed to verify that furimazine remained in the docking active site, using the best docked pose from Vina as the starting structure. OpenFF's SMIRNOFF force field was used to parameterize furimazine, and Amber's ff14SB force field was used to parameterize the protein. Solvation and charge neutralization, minimization, NPT equilibration, and the 100 ns simulations were all carried out with OpenMM. Chimera was then used to identify amino acid residues within 5Åof the consensus docking poses. For the large active site, all residues within 7Åwere selected. These amino acids were presumed to comprise the active site of the protein.

###### B.1.4 NanoLuc Heat Treatment, Solubility Assessment, and Activity Assay

The NanoLuc proteins were expressed as 3 biological replicates of 40 $\mu$L CFPS reactions for each mutant in parallel. Reactions were incubated for 3 hours at 37° C and 280 RPM. Protein reaction replicates were combined, sampled in triplicate for scintillation counting yield and redivided into five aliquots in PCR tubes, where each aliquot was heat treated at one of five temperatures: 37, 50, 57, 63, and 70 degrees C. Heat treatment was conducted for 15 minutes in a PCR thermocycler chamber, or an incubator for the 37° C heat treatment. After heat treatment, all samples were centrifuged for 15 min at 16100 g at 4° C and the supernatant was sampled in triplicate for scintillation counting to assess soluble NanoLuc protein remaining after heat treatment.

#### B.2 WW synthesis and assays

###### B.2.1 Synthesis

WW, BayesDesign and ProteinMPNN peptides were synthesized as C-terminal acids, by microwave-assisted solid-phase peptide synthesis, using a standard Fmoc N$\alpha$ protection strategy. Amino acids were activated by 2-(1H-benzotriazole-1-yl)-1,1,3,3-tetramethyluronium hexafluorophosphate (HBTU, purchased from Advanced ChemTech) and N-hydroxybenzotriazole hydrate (HOBt, purchased from Advanced ChemTech). Fmoc-Gly-loaded Novasyn Wang resin and all Fmoc-protected $\alpha$-amino acids with acid-labile side-chain protecting groups were purchased from CombiBlocks. Peptide variants were synthesized on a 50 $\mu$mol scale. Acid-labile side-chain protecting groups were globally removed and proteins were cleaved from the resin by stirring the resin for 4h in a solution of phenol (0.125 g), water (125 $\mu$L), thioanisole (125 $\mu$L), ethanedithiol 62.5 $\mu$L) and triisopropylsilane (25 $\mu$L) in trifluoroacetic acid (TFA, 2 mL). Following the cleavage reaction, the TFA solution was drained from the resin, the resin was rinsed with additional TFA. Proteins were precipitated from the concentrated TFA solution by addition of diethyl ether ( 40 mL). Following centrifugation, the ether was decanted, and the pellet was dissolved in 40mL 1:1 H2O/MeCN, frozen and lyophilized to remove volatile impurities. The resulting powder was stored at -20° C until purification.

##### B.2.2 Purification and Characterization

Immediately prior to purification, the crude protein was dissolved in 1:1 H2O/MeCN. Proteins were purified by preparative reverse-phase HPLC on a C18 column using a linear gradient of water in acetonitrile with 0.1% v/v TFA. HPLC fractions containing the desired protein product were pooled, frozen, and lyophilized. Proteins were identified by electrospray ionization time of flight mass spectrometry (ESI-TOF, values shown in table 3 below), and purity was analyzed by Analytical HPLC (see Figure 10).

##### B.2.3 Circular Dichroism Spectropolarimetry

Measurements were made with an Aviv 420 Circular Dichroism Spectropolarimeter, using quartz cuvettes with a path length of 0.1 cm. Protein solutions were prepared in 20 mM sodium phosphate buffer, pH 7, and protein concentrations were determined spectroscopically based on tyrosine and tryptophan absorbance at 280 nm in 6 M guanidine hydrochloride + 20 mM sodium phosphate ($\epsilon$Trp = 5690 M-1 cm-1 , $\epsilon$Tyr = 1280 M-1 cm1 ). CD spectra of 50 $\mu$M solutions were obtained from 260 to 200 nm at 25° C and 95° C.

#### C Proofs

**Proposition 1.** *If a sequence s maximizes stability, then it maximizes the probability ratio between the folded and unfolded states.*

*Proof:* Consider two sequences $s$ and $s'$, and assume

$$G(Z'|\text{seq} = s') - G(X'|\text{seq} = s') < G(Z|\text{seq} = s) - G(X|\text{seq} = s).$$

Note that $e^{-x}$ is a monotonically decreasing function, and for monotonically decreasing functions, $a < b \implies f(a) > f(b)$. Thus:

$$e^{-(G(Z'|\text{seq}=s')-G(X'|\text{seq}=s'))} > e^{-(G(Z|\text{seq}=s)-G(X|\text{seq}=s))}$$
$$\frac{p(\text{structure} = Z'|\text{seq} = s')}{p(\text{structure} = X'|\text{seq} = s')} > \frac{p(\text{structure} = Z|\text{seq} = s)}{p(\text{structure} = X|\text{seq} = s)}$$

Since both sides of the inequality are positive, taking the reciprocal of both sides flips the inequality.

$$\frac{p(\text{structure} = X'|\text{seq} = s')}{p(\text{structure} = Z'|\text{seq} = s')} < \frac{p(\text{structure} = X|\text{seq} = s)}{p(\text{structure} = Z|\text{seq} = s)}$$

□

### References

[Bundy and Swartz, 2010] Bundy, B. C. and Swartz, J. R. (2010). Site-specific incorporation of p-propargyloxyphenylalanine in a cell-free environment for direct protein-protein click conjugation. *Bioconjugate Chemistry*, 21(2):255–263. PMID: 20099875.

[Hunt et al., 2022] Hunt, J. P., Zhao, E. L., Free, T. J., Soltani, M., Warr, C. A., Benedict, A. B., Takahashi, M. K., Griffitts, J. S., Pitt, W. G., and Bundy, B. C. (2022). Towards detection of sars-cov-2 rna in human saliva: A paper-based cell-free toehold switch biosensor with a visual bioluminescent output. *New Biotechnology*, 66:53–60.

[Jewett and Swartz, 2004] Jewett, M. C. and Swartz, J. R. (2004). Mimicking the escherichia coli cytoplasmic environment activates long-lived and efficient cell-free protein synthesis. *Biotechnology and Bioengineering*, 86(1):19–26.

[Tomabechi et al., 2016] Tomabechi, Y., Hosoya, T., Ehara, H., ichi Sekine, S., Shirouzu, M., and Inouye, S. (2016). Crystal structure of nanokaz: The mutated 19 kda component of oplophorus luciferase catalyzing the bioluminescent reaction with coelenterazine. *Biochemical and Biophysical Research Communications*, 470(1):88–93.
